# Extensive genomic divergence among 61 strains of *Chlamydia psittaci*

**DOI:** 10.1101/2022.11.10.515926

**Authors:** Konrad Sachse, Martin Hölzer, Fabien Vorimore, Lisa-Marie Barf, Kevin Lamkiewicz, Carsten Sachse, Karine Laroucau, Manja Marz

## Abstract

*Chlamydia (C.) psittaci*, the causative agent of avian chlamydiosis and human psittacosis, is a genetically heterogeneous species. Its broad host range includes parrots and many other birds, but occasionally also humans (via zoonotic transmission), ruminants, horses, swine and rodents. To assess whether there are genetic markers associated with host tropism we comparatively analyzed whole-genome sequences of 61 *C. psittaci* strains. Initially, poorly assembled genomes in public databases were subjected to clean-up, reassembly and polishing. Multiple sequence alignment of the genome sequences revealed four major clades within this species. Clade 1 represents the most recent lineage comprising 40/61 strains and contains 9/10 of the psittacine strains, including type strain 6BC, and 10/13 of human isolates. Clades 2-4 carry strains from different non-psittacine hosts. We found that clade membership correlates with classification schemes based on SNP types, *ompA* genotypes, multilocus sequence types as well as plasticity zone (PZ) structure and host preference. Genome analysis also revealed that i) sequence variation in the major outer membrane porin OmpA can result in 3D structural changes of immunogenic domains, ii) past host change of Clade 3 and 4 strains could be associated with loss of MAC/perforin in the PZ, rather than the large cytotoxin, iii) the distinct phylogeny of atypical strains (Clades 3 and 4) is also reflected in their repertoire of inclusion proteins (Inc family) and polymorphic membrane proteins (Pmps). Altogether, our study identified a number of genomic features that can be correlated with the phylogeny and host preference of *C. psittaci* strains.

## 1 Introduction

*Chlamydia (C.) psittaci* is known as the etiological agent of avian chlamydiosis and human psittacosis [1]. Due to its capability of causing systemic infection with acute to chronic course in poultry and pet birds, as well as its worldwide dissemination [2], it is probably the most important veterinary chlamydial pathogen. Besides, the importance of *C. psittaci* as a human pathogen is often underestimated. Although the big outbreaks of “parrot fever” following large shipments of exotic birds from South America to Europe and North America in the period from 1892 to 1929 are now history, the agent still deserves permanent attention. The zoonotic potential of *C. psittaci* is well documented in the literature [3–5]. Typically, individuals or small groups with previous contact to birds are affected, but fulminant manifestations in humans usually occur only when efficacious antimicrobials are not administered in time. The course of the human disease ranges from asymptomatic to flu-like to severe systemic illness, with the latter manifesting as pneumonia, myocarditis, encephalitis or sepsis. Mild symptoms are seen in most individuals infected, but immunocompromised persons are more likely to develop clinical signs. Occasionally, also apparently healthy individuals can be severely affected [6, 7]. In the last two decades, cases of zoonotic transmission were reported from psittacine birds [8], as well as ducks [9, 10], turkeys [11] and mixed domestic poultry [7] as the main sources. In addition, human-to-human transmission was shown to be a relevant infection route in a number of cases [12–16].

Like all chlamydial organisms, *C. psittaci* is an obligate intracellular bacterium distinguished by a biphasic developmental cycle comprising extracellular and intracellular stages. In the course of evolution, the genomes of all *Chlamydia* spp. have undergone vast condensation, which was shown to have resulted from genome streamlining rather than degradation [17]. The relatively small genome size of approximately 1 Mbp implies the absence of essential cellular pathways and, consequently, reliance on host cells for nutrients, such as amino acids, nucleotides and lipids [18, 19]. *Chlamydiae* are assumed to compensate for this deficiency by co-opting suitable cellular pathways [20,21]. As handling of *Chlamydia* spp. using cell culture requires special expertise and their genetic manipulation is much more difficult than of most other bacteria, analysis of whole-genome sequences can be a viable alternative to characterize strains of interest and provide clues to understand pathogenic properties.

A large number of genome assemblies of varying quality from all *Chlamydia* spp. were published in the last decade. We included *C. psittaci* strains whose genomes genomes sequences, including raw data, were available from the NCBI and ENA databases on February 1, 2021 and had less than 50 scaffolds after reassembly using our own procedure described below. Given the steady advance of sequencing technologies and bioinformatics tools, it is not surprising that these genome assemblies differ significantly in quality, e.g. in scaffold numbers between 1 and 851 and N50 values between 1 Mbp (genome size) and 725 bp. In addition, the quality of gene annotations depends on the quality of the underlying assembly, the used annotation approach, software versions and parameters, as well as the reference database, which still represents a bioinformatics bottleneck even when high-quality genome assemblies are available [22]. These deficiencies in genome quality and annotation status can seriously hamper comparative studies.

Several genomic studies dealt with the comparison of whole-genome sequences among the major *Chlamydia* species [23–27]. A few others addressed questions of genomic diversity of the human pathogens *C. trachomatis* [28] and *C. pneumoniae* [29], as well as the zoonotic agent *C. abortus* [30]. Concerning *C. psittaci*, Read et al. [31] analyzed the genomes of 20 strains from different hosts to suggest events of host switching and recombination along the timeline of phylogenetic evolution. In view of the considerable variation in terms of host preference, growth characteristics and pathogenicity observed among *C. psittaci* strains it seems necessary to study a larger number of field strains to obtain data on intra-species genetic variation at genome level. In a recent comparative study comprising 33 strains of 12 different *Chlamydia* spp., among them 10 strains of *C. psittaci*, we singled out genomic features characteristic for *C. psittaci*, i.e. (i) a relatively short plasticity zone (PZ); (ii) an inclusion membrane protein (Inc) set comprising IncA, B, C, V, X, Y; (iii) the largest chlamydial SinC protein sized 502 amino acids; and (iv) an elevated number of subtype G Pmp proteins (*n* = 14) [26].

In the present paper, we report the findings of a comparative analysis of 61 *C. psittaci* genomes that were deposited in public databases and met our quality requirements. Our study aimed at elucidating the extent of intra-species genomic divergence and searching for possible correlations between genomic and phenotypic parameters.

## 2 Results

### Improvement of genome assemblies and annotation

By February 1, 2021, 71 genome assemblies of *C. psittaci* strains had been uploaded to the NCBI and ENA databases. To ascertain data consistency and comparability, we removed any duplicates and finally included only whole-genome sequences fulfilling the quality criteria stated below. We re-assembled raw sequencing data available for 38 *C. psittaci* strains to achieve a better genome and annotation quality. After all cleaning steps, 11 re-assemblies achieved a better quality and were used instead of the original NCBI genomes in our final genome collection (Table S1). We set the maximum number of scaffolds to be tolerated to 50 to ensure consistency among the finally used assemblies, e.g., for detection of the PZ. In addition, we checked assembly metrics such as the N50 value and the number of potentially fragmented ORFs (via so-called `IDEEL` plots, as described in Stewart et al. [32]) to finally include genome assemblies of 61 strains in this study. Among them, 37 had completely assembled genome sequences (one sequence), 20 consisted of 2–8 scaffolds and another four genomes had 16, 19, 28 or 44 scaffolds, respectively (Table S1). This is a considerable improvement compared to the assembly states appearing in the respective NCBI and ENA entries. The reassembled genomes are deposited in the OSF repository: https://doi.org/10.17605/OSF.IO/RBCA9. In Supplement Figure S1, we compare all re-assembled and original genomes regarding their genome contiguity and N50 values, while the `IDEEL` plots in the OSF repository show the improvement regarding fragmented ORFs. Moreover, re-annotation of all genomes using recent software versions and reference databases helped to reduce the proportion of nonannotated genome features (hypothetical proteins) to 25 % (average 251 of 994 CDS, see Table S1).

### Genome size and core genome

The average genome size of the 61 *C. psittaci* strains was 1 166 132 bp. Strains 99DC5 and WS-RT-E30 were found to harbor the largest (1 175 249 bp) and smallest (1 140 789 bp) genomes of the present panel, respectively. Genome sizes of all strains are provided in Table S1. While the pan-genome was composed of 1 126 CDS, the core genome of this strain panel was calculated to be 904 common CDS (using `RIBAP`), which is 90.9 % of the average 994 CDS detected. The complete output of `RIBAP` is available at the OSF: https://doi.org/10.17605/OSF.IO/RBCA9

### Phylogenetic analysis

To explore the phylogenetic relationship among the strains, we reconstructed a tree based on the alignment of whole-genome sequences (Figure 1A). Our data shows that the species comprises four major clades. The largest clade, consisting of 40/61 strains, includes the type strain 6BC and will be referred to as Clade 1. All psittacine isolates are on this clade alongside some others from humans, cattle, sheep and horses. Genome sequences of Clade 1 strains tend to be highly similar despite the presence of up to six major recombination sites (Figure 1D/E, sites are marked gray). Since typing parameters of these strains are exclusively mainstream they are regarded as typical *C. psittaci* strains.

**Figure 1:**
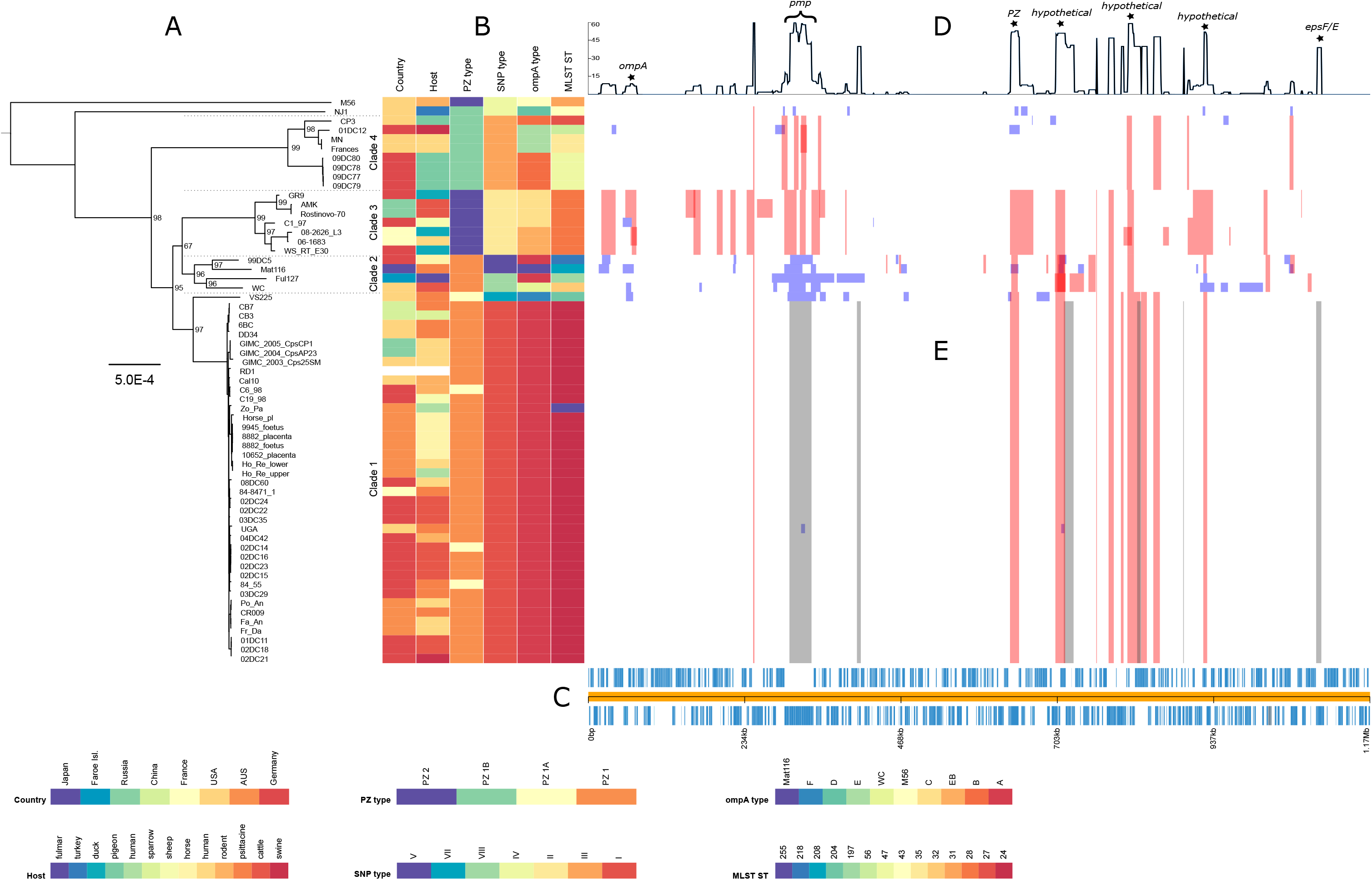
Global phylogeny based on the whole-genome alignment and recombination landscape of 61 *C. psittaci* genomes. The phylogeny (A), with associated metadata (B) is displayed alongside the linearized chromosome (C). Line graph (D) shows the number of recombination events affecting individual genes. Colored blocks (E) indicate inferred recombination events with blue blocks unique to a single isolate and red blocks shared by multiple strains through common descent. Grey blocks represent major recombination sites. Gene annotations are based on type strain 6BC (CP002549.1). The scale bar of the phylogenetic tree corresponds to 5 × 10^−4^ substitutions per nucleotide site and bootstrap values indicate stability of the branches based on 1 000 replicates.

Clade 2 contains only four strains (99DC5, Ful127, Mat116, and WC), which belong to rare SNP and multilocus sequence types, but are otherwise similar to Clade 1 strains. Each of these genomes harbors an unusually high number of unique sequences due to recombination events (blue boxes in Figure 1E).

Clade 3 has seven strains isolated from non-psittacine hosts, such as duck, sheep, cattle and human: (06-1683, 08_2626_L3, AMK, C1/97, GR9, WS-RT-E30, and Rostinovo-70). Their typing parameters are largely atypical, i.e. PZ type 2, SNP type II, *ompA* types EB or C and MLST 28 (Table S1).

Clade 4 is formed by eight strains from non-psittacine hosts, mainly pigeons (MN, 09DC77, 09DC78, 09DC79, 09DC80, Frances, CP3, and 01DC12).

Finally, strains M56 from a muskrat and NJ1 from a turkey are encountered outside the clades, which is consistent with their atypical classification as SNP type IV, ST31 and 43, as well as *ompA* genotypes M56 and D, respectively.

We additionally constructed alternative phylogenetic trees based on (a) the 904 core genes that emerged from `RIBAP`, calculated by `RAxML` (Figure S2), (b) the SNP analysis calculated by `RAxML` (Figure S3) (c) the extracted PZ of 53 strains (Figure S4), and (d) multiple sequence alignment of all 61 OmpA proteins (Figure S5). The assignment of strains to a particular clade in our phylogenetic tree is largely concordant with the classification schemes derived from SNP typing and MLST analysis, as well as *ompA* genotyping and the newly introduced PZ types (see Table S1).

These relationships will be discussed in more detail below.

### Comparative analysis of the plasticity zone

The PZ was defined as the segment flanked by genes *accB* (5’) and *guaB* (3’). A non-fragmented PZ could be extracted from genome sequences of 53 strains. The zone varied in size from 22 534 nt (WS-RT-E30) to 30 180 nt (C6/98). In the genome of strain 6BC, the PZ is located between positions 624 296 and 653 440. The major ORFs are compiled in Table 1.

**Table 1:** Major open reading frames in the plasticity zone of *C. psittaci* strain 6BC. Len – length [nt]; Dir – direction; + − forward, plus strand; - – reverse, minus strand;

| No | Name | Start | End | Len | Dir | Comment |
| --- | --- | --- | --- | --- | --- | --- |
| 1 | *accB* | 1 | 501 | 501 | + | Biotin carboxyl carrier protein of acetyl-CoA carboxylase |
| 2 | *accC* | 501 | 1856 | 1356 | + | Biotin carboxylase (also Biotin acetyl-CoA carboxylase subunit) |
| 3 | DUF648 domain-containing protein_1 | 2008 | 3438 | 1431 | + | Domain of Unknown Function |
| 4 | DUF648 domain-containing protein_2 | 3558 | 5132 | 1575 | + | Domain of Unknown Function |
| 5 | Hypoth. protein ORF 141 | 5340 | 6296 | 957 | - |  |
| 6 | Hypoth. protein ORF 137 | 6664 | 7605 | 942 | - |  |
| 7 | MAC/perforin family protein | 7794 | 8477 | 684 | - | possibly includes phospholipase D domain |
| 8 | DUF1389 domain-containing protein | 8853 | 9950 | 1098 | - | Domain of Unknown Function |
| 9 | putative membrane protein | 9880 | 10398 | 519 | - |  |
| 10 | *toxB* (LifA/Efa1-related large cytotoxin) | 10434 | 20507 | 10074 | - | also: Lymphostatin (in E.coli), Cysteine protease, YopT-type domain protein, TcdB toxin N-terminal helical domain protein |
| 11 | Hypoth. protein ORF 67 | 20788 | 21549 | 762 | + |  |
| 12 | MAC/perforin | 21962 | 24430 | 2469 | + | Membrane-attack complex/perforin (MACPF) superfamily protein |
| 13 | Hypothet. protein G50_0603 | 24539 | 25060 | 522 | - |  |
| 14 | ADA | 25124 | 26524 | 1401 | - | Adenosine/AMP deaminase |
| 15 | *guaA* | 26515 | 28053 | 1539 | - | GMP synthase |
| 16 | *guaB* | 28068 | 29144 | 1077 | - | Inosine-5'-monophosphate dehydrogenase |

Comparison of PZ structures based on multiple alignment of 53 sequences revealed four different types, which we define as types 1, 1A, 1B and 2, as depicted in Figure 2 (sequence similarity values in Table S2). In PZ type 1, which was encountered in 32 strains, the complete set of 16 ORFs was identified. The main elements include the 5’-terminal biotin modification operon (*accB*, *accC*), the large cytotoxin (*toxB*) and MAC/perforin in the central region, as well as the purine synthesis and recycling operon (*ADA*, *guaA, guaB)* at the 3’ terminus. PZ type 1A is distinguished by a fragmented or disrupted cytotoxin, e.g. in strain VS225 with four smaller proteins instead of one large molecule, also in strains C6/98, 02DC14, and 84-55 (three fragments). Type 1B can be recognized by the fragmented MAC/perforin (strains MN, 09DC77, 09DC78, 09DC79, 09DC80, Frances, CP3, 01DC12, NJ1). These three PZ types share a highly homologous structure with overall nucleotide sequence identity values above 95 %. PZ type 2, which was encountered in all seven strains of Clade 3 (06-1683, 08-2626_L3, AMK, C1/97, GR9, Rostinovo-70, WS-RT-E30) and M56, represents a reduced PZ version and is characterized by the absence of six ORFs at the 3’-end (nos. 11-16 in Table 1 and Figure 2) including MAC/perforin and the purine synthesis operon. The PZ of this type is about 20 % shorter than in the other strains.

**Figure 2:**
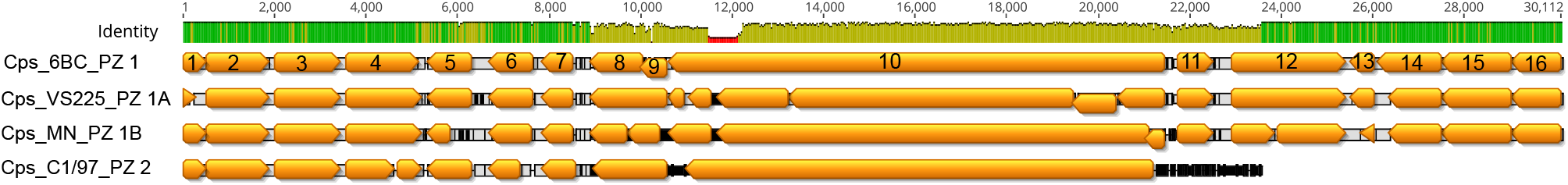
Alignment of the plasticity zones of strains 6BC, VS225, MN and C1/97 representing PZ types 1, 1A, 1B, and 2, respectively. Numbering of ORFs as in Table 1.

In eight strains, i.e. Fa_An, Fr_Da, CB3, CB7, 9945_foetus, 8882_placenta, 8882_foetus, and 10652_placenta, PZ elements were located on separate scaffolds, thus precluding extraction of a contiguous PZ. As these strains could not be included in the multiple sequence alignment, the PZ sequence of strain 6BC was mapped to each of these genome sequences. As a result, the presence of all major CDS, including the terminal *accB* and *guaB*, the large cytotoxin and MAC/perforin genes were confirmed, which indicates a type 1 PZ in these strains.

### The family of polymorphic membrane proteins (Pmps)

To explore the spectrum of Pmp family members present in the 61 strains, we conducted multiple protein blast analyses using the known Pmp sequences of strain 6BC as queries. The results confirm that all 21 family members are present in all 61 genomes (Table S3). While strains of Clade 1 were found to carry Pmps that are often identical and generally highly similar to their equivalent in strain 6BC, sequence similarity with the rest of the strains was markedly lower. The data in Table 2, which shows representative strains, reveal that the lowest identity percentages were seen in Clade 3 and the outlying strain M56. In addition, some of the Pmps of Clade 4 strains tended to align only partially to the 6BC equivalents, which could mean that they are shorter or contain distinct sequence elements. Regarding individual Pmps, there is considerable variation among strains in some genomic loci, notably pmp12-15 and pmp17-21, all of which belong to subtype G/I. Pmp15 and Pmp17 are the most variable family members with the average amino acid (aa) identity to the 6BC equivalents being only 75,90 %. On the other hand, Pmps1-11 as well as 16 and 22 showed less sequence variation with average aa identity values above 95 % (Table S4).

**Table 2:** Amino acid sequence divergence of Pmp repertoires among *C. psittaci* strains from different phylogenetic clades.

| Strain: | 02DC15 (Clade 1) |  | Mat116 (Clade 2) |  | C1/97 (Clade 3) |  | CP3 (Clade 4) |  | NJ1 (outside) |  | M56 (outside) |  |
| --- | --- | --- | --- | --- | --- | --- | --- | --- | --- | --- | --- | --- |
| Pmp [no._subtype] | % ident | a/q | % ident | a/q | % ident | a/q | % ident | a/q | % ident | a/q | % ident | a/q |
| Pmp1_B | 100.00 | 1.00 | 98.72 | 1.00 | 99.29 | 1.00 | 97.77 | 1.00 | 94.56 | 1.00 | 87.42 | 1.00 |
| Pmp2_A | 100.00 | 1.00 | 99.79 | 1.00 | 99.89 | 1.00 | 99.68 | 1.00 | 99.57 | 1.00 | 96.68 | 1.00 |
| Pmp3_E/F | 100.00 | 1.00 | 93.41 | 1.00 | 86.50 | 1.00 | 86.10 | 1.00 | 95.10 | 1.00 | 70.69 | 1.00 |
| Pmp4_E/F | 99.90 | 1.00 | 82.12 | 0.66 | 94.49 | 1.00 | 96.88 | 1.00 | 94.07 | 1.00 | 72.33 | 1.00 |
| Pmp5_E/F | 100.00 | 1.00 | 100.00 | 1.00 | 99.44 | 1.00 | 99.15 | 1.00 | 96.61 | 1.00 | 65.73 | 1.01 |
| Pmp6_H | 100.00 | 1.00 | 94.01 | 0.75 | 94.92 | 1.00 | 94.82 | 1.00 | 93.60 | 1.00 | 89.95 | 1.00 |
| Pmp7_G/I | 100.00 | 1.00 | 92.56 | 1.01 | 93.38 | 1.00 | 91.47 | 1.00 | 91.66 | 1.00 | 86.39 | 1.00 |
| Pmp8_G/I | 100.00 | 0.80 | 99.74 | 0.90 | 99.65 | 1.00 | 97.71 | 0.46 | 95.77 | 1.00 | 84.43 | 0.81 |
| Pmp9_G/I | 100.00 | 0.75 | 89.10 | 0.46 | 93.02 | 1.00 | 97.82 | 0.34 | 79.59 | 0.77 | 68.05 | 0.48 |
| Pmp10_G/I | 100.00 | 1.00 | 99.64 | 1.00 | 99.26 | 0.64 | 98.89 | 0.64 | 97.14 | 1.00 | 91.91 | 1.00 |
| Pmp11_G/I | 100.00 | 1.00 | 99.53 | 0.99 | 99.29 | 1.00 | 99.41 | 1.00 | 96.82 | 1.00 | 90.58 | 1.00 |
| Pmp12_G/I | 100.00 | 1.00 | 84.58 | 1.00 | 77.54 | 1.00 | 84.68 | 0.67 | 85.71 | 1.00 | 81.62 | 1.00 |
| Pmp13_G/I | 99.88 | 1.00 | 78.05 | 1.01 | 74.76 | 1.00 | 77.25 | 0.67 | 77.62 | 1.01 | 84.43 | 1.01 |
| Pmp14_G/I | 100.00 | 1.00 | 79.16 | 1.01 | 77.27 | 1.00 | 81.16 | 0.67 | 79.93 | 1.01 | 80.64 | 1.00 |
| Pmp15_G/I | 83.43 | 1.00 | 93.92 | 1.00 | 77.90 | 1.00 | 96.12 | 0.66 | 86.45 | 1.00 | 80.12 | 1.00 |
| Pmp16_G/I | 100.00 | 1.00 | 99.78 | 1.00 | 99.57 | 1.00 | 98.71 | 1.01 | 98.38 | 1.00 | 90.82 | 1.00 |
| Pmp17_G/I | 83.43 | 1.00 | 93.92 | 1.00 | 77.90 | 1.00 | 96.12 | 0.66 | 86.45 | 1.00 | 80.12 | 1.00 |
| Pmp19_G/I | 77.66 | 1.02 | 77.00 | 1.02 | 99.40 | 1.00 | 80.42 | 0.68 | 80.12 | 1.01 | 80.83 | 1.01 |
| Pmp20_G/I | 99.88 | 1.00 | 81.34 | 1.00 | 77.09 | 1.00 | 83.63 | 0.67 | 82.63 | 1.00 | 80.02 | 1.00 |
| Pmp21_G/I | 84.68 | 1.01 | 86.64 | 1.01 | 79.34 | 1.00 | 84.30 | 0.67 | 90.12 | 1.00 | 88.92 | 1.00 |
| Pmp22_D | 100.00 | 1.00 | 99.28 | 1.00 | 87.57 | 1.00 | 99.48 | 0.88 | 97.85 | 1.00 | 99.35 | 1.00 |

% ident Percentage of identities shared with query sequence of *C. psittaci* strain 6BC
a/q Ratio of aligned target sequence length (blast hit) to query sequence length.

### The Inc protein family

At least 11 Inc proteins were identified by the annotation pipeline. Nine of them represent different members of the IncA family, which have been arbitrarily designated IncA family protein 1-9. Five of them were found in all 61 strains, while three others were absent in the seven strains of Clade 3. Major inter-strain sequence variation was observed in three proteins, IncA family proteins 3 and 5 as well as IncV, again with Clade 3 strains standing out. The main results are given in Table 3.

**Table 3:** Presence of Inc proteins in *C. psittaci* strains and variation in atypical strains. IncA family protein (fam prot) 4 occurs only in Clade 1 and 2 (*) and is truncated in Clade 3 and 4. Group numbers refer to `RIBAP` output. Size refers to 6BC [aa]. MinID – Minimum identity among typical strains [%].

| Annotation | Size | No. of strains | MinID | Exceptions |
| --- | --- | --- | --- | --- |
| IncA fam prot 1 (group 358) | 372 | 61 | 97.8 |  |
| IncA fam prot 2 (group 514) | 224 | 61 | 98.7 |  |
| IncA fam prot 3 (group 890) | 352 | 61 | 99.4 | Clade 3 strains with low similarity (75 %) |
| IncA fam prot 4 (group 915) | 461 | 35* | 99.3 | Truncated in strains of Clades 3+4 |
| IncA fam prot 5 (group 939) | 238 | 61 | 98.7 | Clade 3 strains with low similarity (60 %); truncated in Mat116, NJ1. |
| IncA fam prot 6 (group 698) | 382 | 61 | 96.6 |  |
| IncA fam prot 7 (group 886) | 342 | 54 | 99.4 | Missing in Clade 3 strains |
| IncA fam prot 8 (group 931) | 479 | 54 | 99.4 | Missing in Clade 3 strains |
| IncA fam prot 9 (group 923) | 810 | 52+2 | 99.4 | Missing in Clade 3 strains (also found in strains 8882_placenta and 8882_foetus via mapping) |
| IncB (group 404) | 202 | 61 | 97.5 |  |
| IncV (group 898) | 383 | 61 | 95.8 | Clade 3 strains with low similarity (85 %) |

### Variation of OmpA sequences in *C. psittaci* strains

An alignment of extracted OmpA sequences (`RIBAP` group 879) confirmed that all 61 strains are equipped with this outer membrane porin. While the complete protein of 402 aa was seen in the vast majority of strains, a group of seven strains harbored slightly shortened versions: GR9, C1/97, AMK, Rostinovo-70 (all from Clade 3), VS225 (Clade 1), WC (Clade 2) and NJ1 (outside). OmpA molecules in that group have lost up to 11 aa, thus rendering sequence identity to the 6BC counterpart as low as 82.84 %. The range of OmpA sequence diversity among the present 61 strains is illustrated in a `RAxML` tree in Figure S5.

To better understand the consequences of the observed sequence variation in the framework of the OmpA porin 3D structure, we compared predicted 3D OmpA structures from two “antagonist” strains, 6BC (typical strain) and C1/97 (atypical strain), using `AlphaFold` [33]. Overall, OmpA/6BC is an all-β fold protein with an accessory N-terminal α-helix (aa positions 1-22) representing the signal peptide. The β-barrel of the transmembrane porin fold resides in the bacterial membrane from which the extra-membrane region emanates (Figure 3A). The core of this region is formed by a short three-stranded β-sheet (159-161, 309-312, 352-356), which is surrounded by the solvent-exposed variable domains (VDs) 1, 2, 3 and 4 (Figure 3B). In OmpA/6BC, the VD1 (83-107), VD2 (164-177) and VD4 (317-350) of the antigen contain three pronounced loop structures without clearly assigned secondary structure, whereas VD3 (228-265) comprises a two-stranded β-sheet. As expected, the superposition of the 6BC and C1/97 structures reveals high overall structural agreement, in particular in the transmembrane β-barrel domain as well as in VD3. Notable differences, however, occur in backbone conformations of the extra-membrane VDs 1, 2 and 4 (Figure 3C). At the same time, VD1, VD2 and VD4 represent the least reliable regions of the `AlphaFold` predictions due to the lower overall pLDDT score, which is generally recognized as predictor for structural disorder and dynamic flexibility of the protein structure [34]. In pairwise alignment of 6BC and C1/97 OmpA sequences, the latter shows several deletions in VD1 (4-aa gap in pos. 100-103), VD2 (4 aa missing between 167 and 176) and VD4 (3-aa gap 342-344). In addition, VDs contain notable aa exchanges, i.e. 14 aa in VD1, 6 in VD2 and 12 in VD4. Remarkably, these strain sequence variations are limited to solvent-exposed regions of VD1, VD2 and VD4, but have no effect on the overall structural porin scaffold.

**Figure 3:**
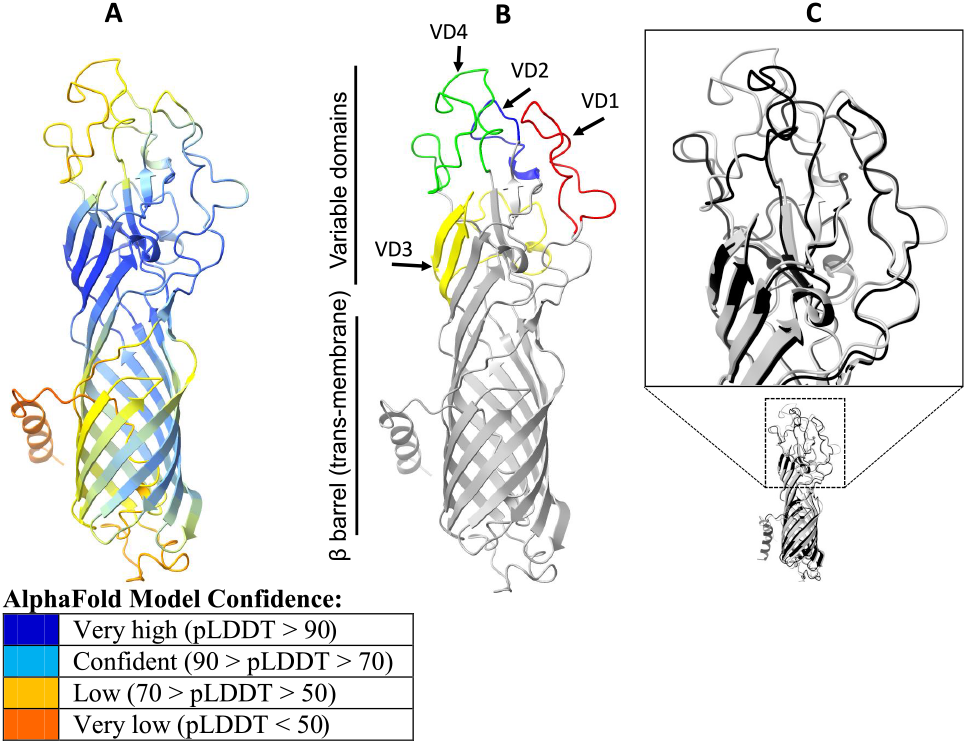
Predicted 3D structure of OmpA from *C. psittaci* strain 6BC and comparison with strain C1/97. **A** Structure prediction of OmpA from strain 6BC using `AlphaFold`. Color code indicates prediction reliability based on perresidue confidence score (pLDDT). **B** Highlighted locations of variable domains (VD) 1 to 4 in strain 6BC. **C** Superposition of predicted OmpA structures in strains 6BC (gray) and C1/97 (black) with zoomed inset of the VD regions. The 3D structures were rendered using `ChimeraX`.

### Other potentially important loci

To explore the stability of presumably non-variable genomic loci that are encoding potential virulence factors, we analyzed the multiple sequence alignments provided in the respective RIBAP groups. The findings are summarized in Table 4. None of the seven loci proved 100 % identical in all strains. While the CADD and FtsW proteins were found to have only a few variable positions, there was more heterogeneity in the other loci, e.g. SinC, where the strains of Clade 3 harbored a protein variant of lower homology. It is also noteworthy that sequences of the histone-like protein pair HctA/B differed among a number of strains, and HctB was absent in four *C. psittaci* strains, a feature shared with some strains of *C. avium* and *C. gallinacea* [26].

**Table 4:** Presence of potential virulence factors in *C. psittaci* strains and variation in atypical strains. Groups numbers refer to RIBAP output groups. Size refers to 6BC [aa]. MinID – Minimum identity among typical strains [%].

| Gene product | Size | No. of strains | MinID | Exceptions |
| --- | --- | --- | --- | --- |
| CADD (group 207) | 245 | 61 | 99 |  |
| CPAF (group 684) | 605 | 61 | 95 | M56 only 91.9 % id. to other strains |
| FtsW (group 619) | 384 | 61 | 99 |  |
| HctA (group 702) | 123 | 61 | 99 | 117-aa variant in Clade 4 members plus Ful127; M56 only 95.1 % id. |
| HctB (group 867) | 197 | 57 | 99 | 154-aa variant (78.2 % id.) in C19/98, C6/98; missing in CP3, M56, NJ1, VS225 |
| SinC (group 897) | 502 | 61 | 99 | 503-aa variant (86.5 % id.) in Clade 3 members |
| TarP (group 742) | 874 | 61 | 99 | M56 only 88.6–89.1 % id., truncated in Mat116 (826 aa) |

### PZ, SNP, *ompA* and MLST types vs. host preference

The origins and typing results of all 61 strains are given in Tables S1 and S5. To explore a possible relationship between the various genotypes on the one hand and the animal host on the other, we tested for enrichment using Fisher exact test (see Sec. 4). We observed an enrichment of the pigeon host with the PZ type 1B, SNP type III and *ompA* genotype B (*n* = 5, p-value < 1.6 · 10^-7^), as well as of the duck host with PZ type 2, SNP type II and *ompA* genotype EB (*n* = 2, p-value < 0.005). PZ type 1, which is associated with Clade 1, did not present an enrichment for a particular host, but was enriched when grouping psittacine, human and cattle host (*n* = 22, p-value < 0.005).

## 3 Discussion

To ensure accurate annotation and correct genome analysis our initial efforts aimed at improving the assembly status of those whole-genome sequences that were poorly assembled or still in draft state. The limitations posed by incompletely assembled genomes are well known [35, 36], particularly their negative impact on annotation quality [22]. Thus, using state-of-the-art algorithms we decontaminated and re-assembled the raw reads of those chlamydial genomes that were of insufficient quality. Based on this harmonized genome set (validated via `QUAST, Icarus`, and `IDEEL` plots; see Figure S1 and the OSF repository), our subsequent annotation strategy yielded better comparability among all finally included assemblies. To have a closer look at the divergence of specific loci in *C. psittaci*, a refined core gene set was calculated, which also considered orthologous genes of low sequence similarity (≥ 60 %).

### Genomic regions of lower synteny

The **PZ** of *C. psittaci* is medium sized compared to *C. suis* (82 505 nt) or *C. avium* (5 694 nt) [26], and in contrast to *C. caviae, C. felis* and *C. pecorum*, the tryptophan biosynthesis operon is missing [24, 37]. Although variations in size and contents are more extensive among chlamydial species the diversity within *C. psittaci* is still considerable, thus justifying the introduction of four different structural types, i.e. 1, 1A, 1B, and 2. Among the strains examined here, C6/98 had the largest (30 180 nt) and WS-RT-E30 the shortest (22 534 nt) PZ.

The most prominent locus in the PZ is the *toxB* gene encoding the large cytotoxin. The fact that its annotation is still a challenge for most algorithms may be due to its large size (3 358 aa in 6BC, 3 159 aa in C1/97) and/or multifunctional domain content. The *toxB* gene product is an ortholog of lymphostatin/EFA-1, a toxin known from *E. coli* (EPEC and EHEC) and also *Citrobacter rodentium*. This protein carries three enzymatic activities attributed to motifs of glycosyl-transferase (D-X-D), protease (C, H, D) and aminotransferase (TMGKALSASA) [38]. Therefore, the molecule is often annotated as LifA/Efa1-related large cytotoxin [39] or cysteine protease YopT-type domain protein [40]. Additionally, based on homology to *Clostridioides difficile*, the 150-aa segment near the N-terminus carrying glycosyltransferase activity is designated TcdB toxin N-terminal helical domain protein in the UniProt database^1^.

While these features render the large cytotoxin a straightforward candidate for virulence factor, experimental evidence in a chlamydial context has yet to be obtained. Reports showing clostridial cytotoxins being capable of Ras superfamily inactivation [41, 42] and host cytoskeleton disassembly [40, 43] could be an incentive to pursue this path. In the present study, we observed truncated versions of the *toxB* gene in strains VS225, C6/98, 02DC14, and 84/55 classified as PZ type 1A. However, disruption of this gene did not coincide with different host tropism or other obvious distinctions compared to strains harboring the full-size *toxB*. This is contrasting our findings on strains carrying genes encoding another PZ protein that contains a membrane attack complex/perforin (MACPF) domain. These immune effectors are part of eukaryotic defense mechanisms and can induce cell killing through targeting microbial or host membranes. In the course of co-evolution with the host, chlamydiae have acquired their own MACPF-domain protein. It is assumed that this has enabled chlamydial organisms to partially resist MACPF effector mechanisms from the host and to facilitate their own infection [44]. The orthologous MACPF-domain protein of *C. trachomatis* was suggested to be able to permeabilize the inclusion membrane [45]. We found that all genomes having a type 1 or 1A PZ harbored an ORF encoding MACPF, which was sized 2 469 nt in strain 6BC (ORF12 in Figure 2). The translated protein of 822 aa has a phospholipase D domain in its N-terminal region, however without homologs outside the *Chlamydia* spp. A recent study also showed differential expression of the gene in cell culture [46]. In addition, another ORF in the PZ was annotated as MAC/perforin family protein (ORF8 in Figure 2) in a number of strains. Our finding that presence or absence of the intact MACPF gene coincides with host tropism underlines the importance of this factor and will be discussed below.

The data obtained in this study raises important questions on the functionality of MACPF as a potential virulence factor, as well as the annotation and identity of the ORF8 product, all of which should be addressed in future laboratory studies.

The **Pmp family** consists of autotransporters with surface-exposed and membrane-translocated domains. Members of this highly variable and complex protein family are regarded as virulence factors [47] and/or adhesins and immune modulators [48].

In the present study, Pmps were annotated as autotransporter domain-containing proteins. It was known from previous genome studies that strain 6BC possessed 21 different Pmps [26, 47], and the study by Wolff et al. [49] provided some insight into *pmp* locus divergence among 12 *C. psittaci* strains. These authors found that subtype G Pmps had the highest degree of divergence in *C. psittaci* genomes. The same observation was made later in an analysis of *C. abortus, C. avium, C. caviae, C. felis, C. gallinacea*, and *C. pecorum* genomes [26].

The present data indicate that eight of 14 subtype G/I Pmps, are subject to considerable strain-to-strain sequence divergence, i.e. Pmps 12-15, 17 and 19-21 (Table 2). As can be seen from Figure S6, these variable loci are arranged in two clusters located in separate genomic regions, from position 319 411 to 335 394 and from 707 548 to 715 415, respectively. In contrast, Pmps 1, 2 and 22 (subtypes B, A and D), which are located outside the two variable genome clusters, represent the most stable family members. While amino acid identity values among Pmp2 sequences ranged from 96.7 to 100 % (compared to the 6BC reference), Pmp17 varied from 40.9. to 100% among all strains (Table S4). Interestingly, Pmp17 was suggested to be a key player in host adaptation [50]. Given the extent of strain-to-strain sequence variation observed with some of the G/I subtype Pmps it is important to note that we found representatives of all 21 family members in all 61 strains. In addition to the complete set of 21 Pmps, our multiple protein blast analysis identified a larger number of possibly truncated low-similarity hits in all strains. The significance of these Pmp-like items is unknown and will require future studies.

The family of **inclusion membrane (Inc) proteins** utilizes approximately 4% of the coding capacity in chlamydial genomes [51] and is rather heterogeneous in terms of sequence similarity, which represents a real challenge to annotation tools. At the same time, Incs share a common structural feature: They are inserted in the inclusion membrane via type III secretion. If exposed to the cytosol, some of them are among the major immunogens of *Chlamydia* spp. [27, 52]. Therefore, strain-to-strain differences in Inc protein sequences could result in different pathogenic properties and host tropism.

In analogy to the nomenclature in *C. trachomatis*, the presence of six Inc protein family subtypes was suggested in *C. psittaci*, IncA, B, C, V, X, and Y, the latter three only provisionally assigned [26]. Our analysis revealed 11 individual Incs, which is probably only the tip of the iceberg, since up to 59 family members have been predicted for *C. trachomatis* and 92 for *C. pneumoniae* [51]. The main characteristics of the items identified here are i) highly conserved sequences of all identified Incs among the typical strains (Clades 1, 2 and partly 4), ii) low-similarity Inc variants in four strains, all belonging to Clade 3, and iii) three individual Incs absent in Clade 3 strains. The finding of Clade 3 carrying markedly distinct sets of Inc proteins is remarkable because a study of *C. trachomatis* suggested that some of the more divergent Incs clustered according to clinical groupings and could contribute to tissue tropism [53].

### Outer membrane porin (OmpA)

The *ompA* gene was shown to have experienced the highest rate of recombination in the *C. trachomatis* genome [28]. Structural variation in the OmpA antigen, also called major outer membrane protein (MOMP), was the basis of *C. psittaci* serotyping introduced in the 1990s [54]. Later on, the corresponding *ompA* genotypes were defined and became a more practicable equivalent to serotypes [55]. Previous attempts of correlating *ompA* genotypes with host preference revealed tendencies, but remained tentative [4]. Due to high sequence variation among *C. psittaci* strains (see Figure 1B), the *ompA* locus is often missed as a core gene. However, based on our analysis, which also considers genes of lower sequence similarity, we were able to reconstruct a core genome including *ompA* (see `RIBAP` group 879 in the OSF supplement). The fact that the `RAxML` tree inferred from OmpA protein sequence alignment of all 61 strains (Figure S5) shows the same grouping of strains as in the trees based on genomes and PZ, indicates that this locus could also be used as a marker of host tropism.

Since the well-known OmpA sequence variations among *C. psittaci* strains have not yet been addressed at the 3D structural level, we compared the protein structures of two antagonist strains, 6BC (Clade 1) and C1/97 (Clade 3). They belong to different *ompA* genotypes, A and C, and originate from different hosts, parrot and sheep, respectively. Our *in silico* analysis was facilitated by the use of `AlphaFold`, a new AI system predicting 3D protein structures with high accuracy [33, 56]. While the predicted structures exhibit the expected hallmarks of a porin, it also reveals strain-to-strain differences in the loops formed by VDs 1, 2 and 4. The presence of flexible VD1, VD2 and VD4 loops in OmpA and the strain-to-strain sequence variations are likely a result of adaptation due to interaction with complementary protein surfaces. The analyzed 3D structure and sequence properties of VD1, VD2 and VD4 provide a rationale to the required structural plasticity when the antigen is facing antibodies from different host immune responses. Thus we were able to show that 3D structural differences between OmpAs from strains of different origin are indeed detectable.

While this is the first study to visualize 3D OmpA structural variations, we are aware that it is still a singular finding. Nonetheless, the observation could be potentially important for the study of host-pathogen interaction. Verification of this intriguing speculation will require systematic investigation including wet-lab experiments.

### Phylogeny and host tropism

Although the topic of host preference of *C. psittaci* was addressed in previous studies [26, 49], many questions have remained unanswered to date. As a general conclusion, a yet incompletely known panel of gene products rather than a single factor seems to determine this property of chlamydial strains.

In the present study, we observed a remarkable similarity in the topology of different phylogenetic trees reconstructed from genomic data, i.e. whole genomes, OmpA, core genes, SNPs, and PZ (Figures 1A, S2, S3, S4, S5). With a few exceptions, membership of the major clades or lineages proved stable among these trees, thus indicating that all five datasets reflect phylogeny.

As only whole-genome sequences deposited in public databases could be included in our study, the present choice of *C. psittaci* strains is arbitrary, so that our data cannot comprehensively reveal the geographical distribution and genetic divergence among all naturally occurring strains. Nevertheless, our phylogenetic analysis includes the largest number of *C. psittaci* strains examined so far and allows some interesting insight into the history of this species and the dynamics of its evolution.

The history of individual *Chlamydia* spp. is still poorly understood. Based on analysis of 20 genomes, Read et al. [31] suggested a high evolution rate of 1.68 × 10^-4^ mutation/site/yr (175 SNPs per year) for the species of *C. psittaci*. Their findings are consistent with the assumption that the most recent common ancestor can be dated to the era of the colonization of South America in the 16th to 18th centuries. However, Hadfield et al. [28] argued that this evolution rate is probably an overestimation.

The present genome data revealed the existence of four major clades plus two outlying strains in this species (Figure 1A). It appears straightforward that Clade 1 carrying the majority of strains emerged late on the phylogenetic timeline, possibly as late as the early 20th century [31]. Also the overall genetic homogeneity among these strains suggests recent emergence, which is consistent with reports of large psittacosis outbreaks in the decades before and after 1900. The limited genetic divergence on this clade could also imply that the isolates from non-avian hosts were previously acquired from birds. While originating from 5 different avian (4 psittacine birds and 1 sparrow) and seven non-avian host species including humans, cattle and horses, it is remarkable that 90 % of all psittacine strains are on Clade 1 (another one only on adjacent Clade 2). In analogy to the clade of LGV strains in *C. trachomatis* [28], members of Clade 1 include more virulent strains than the other clades.

The only psittacine strain on Clade 2, Mat116, is from Japan, which could explain its genome being distinct from Clade 1 because of the large geographical distance from most of the other strains.

Clade 3 represents the most genetically distinct lineage of the species. It seems to carry both mammalian and avian strains, however no psittacine strains. In view of the low number of members (*n* = 7) it is not yet certain that psittacine hosts can be excluded, but if so, a switching event from psittacine to non-psittacine host could have initiated the development of this lineage. Notably, the three strains isolated from ducks are encountered on this clade. Ducks are the main *C. psittaci* host among domestic poultry, and these strains seem to be pathogenic to humans as cases of transmission were reported regularly [4, 57–59].

Clade 4 was probably the first to separate from the most recent common ancestor. Similar to members of Clade 3, these eight strains from non-psittacine hosts are distinguished by the loss of MACPF in the PZ, as well as aberrant repertoires of Incs and Pmps.

Overall, the present study provides novel insights into the diversity of the species *C. psittaci*. The findings of this comparative genome analysis will hopefully improve our understanding of possible links between genomic features and phenotypic traits.

## 4 Material and Methods

### Chlamydial strains

We included *C. psittaci* strains whose genome sequences, including raw data, were available from the NCBI or ENA databases on February 1, 2021, and had less than 50 scaffolds after reassembly using our own procedure described below. Strain designations, host organisms and assembly states are given in Table S5. A more detailed overview with basic characteristics and database accession numbers of all 61 *C. psittaci* strains is given in Table S1.

### Reassembly, clean-up and normalization of genome sequences

To reduce technical bias, the same tools and software versions were used to quality-control and *de novo* assemble all single and paired short-read data sets. For paired-end data sets, the respective input parameter settings were adjusted for each tool. First, reads were quality-trimmed using fastp v0.20.1 [60] with parameters `−5 −3 -W 4 -M 20 −l 15 −x −n 5 -z 6`. Subsequently, *de novo* assembly was performed using `SPAdes` v3.14.1 [61] with parameters `--careful --cov-cutoff auto`. Afterwards, the initial assemblies were subjected to two additional polishing rounds using Pilon v1.23 [62]. We used BWA-MEM v0.7.17 [63] to map the quality-controlled short reads to each respective assembly as input for the polishing steps. Finally, we used Bandage v0.8.1 [64] to examine the assembly graphs and their contiguity.

We used the decontamination workflow Clean (v0.2.0)^2^ before and after reassembly to remove foreign sequences from the genomes in public databases, with sequences of *Homo sapiens, Chlorocebus sabeus, Gallus gallus, Macaca nemestrina*, phiX and *C. psittaci* plasmids being considered to be potential contaminants. After reassembly, all strains with more than 50 scaffolds were discarded. We justify this cut-off based on our additional assessment of assembly quality. First, we used `QUAST` v5.2.0 [65] to calculate various metrics between the original and re-assembled genomes and in particular visualized contig N50 and assembly contiguity using the built-in `Icarus` tool [66]. Secondly, to assess ORF quality and compare it in the original and re-assembled genomes, we ran a Snakemake pipeline^3^ implementing an approach called `IDEEL` as previously presented [32]. Briefly, annotated ORFs are translated into proteins and compared against a reference database to plot their completeness. To facilitate genome comparison, whole-genome sequences of all finally included 61 strains were normalized using a custom python script (located as helper script in the `RIBAP` repository^4^) so that the *hemB* (delta-aminolevulinic acid dehydratase) gene appeared in the initial position.

### Annotation and processing of genome sequences

All refined genome sequences were processed using the `Roary` ILP Bacterial Annotation Pipeline (`RIBAP`) v0.6.2^5^, which was recently developed in our group [26]. This pipeline performs annotation, core gene set calculation, alignments and phylogenetic reconstructions of homologous genes in `RIBAP` groups.

As an initial operation, annotation was conducted using `Prokka` v1.14.5 [67] with the `--proteins` option and the annotated GenBank file of *C. psittaci* 6BC (CP002549.1) as reference to ensure consistency in gene denominations. After calculating an initial core gene set with `Roary` v3.13.0 [68], `RIBAP` performs a less stringent all-versus-all `MMSeqs2` v10.6d92c [69] approach to find potential homologous genes missed by `Roary`. Based on all pairwise comparisons, two `Roary` clusters are merged into one `RIBAP` group if the majority shows sufficient homology. Any gene from the core gene set has to be present in all input genomes. The output of a `RIBAP` run includes a searchable and interactive HTML table featuring the final `RIBAP` groups. This includes gene designation, gene description, a color-coded heat map based on `Roary` assignments at different similarity thresholds, as well as a phylogenetic tree based on multiple sequence alignment (MSA) of the CDS contained in the respective `RIBAP` group.

### Whole genome SNP analysis

Each assembly file was processed with the `ART-MountRainier` simulation tool, which generates synthetic paired-end reads with coverage of 50 [70]. These reads were aligned and mapped against the reference sequence of *C. psittaci* 6BC (CP002549.1) using the `BWA` algorithm implemented in `BioNumerics` v7.6.1 (bioMérieux, Applied Maths, Sint-Martens-Latem, Belgium) with minimum 90 % of sequence identity. SNPs were identified using the `BioNumerics` wgSNP module and then filtered using the following conditions: minimum 5 X coverage to call a SNP, removal of positions with at least one ambiguous base, one unreliable base or non-informative SNP and minimum inter-SNP distance of 25 bp.

### Phylogenetic and recombination analyses

A wholegenome alignment (WGA) was performed on the 61 *C. psittaci* genomes using `SKA` v1.0^6^. Prior to that alignment, fragmented genomes with more than one scaffold were mapped on the 6BC reference genome (`NC_015470.1`) using `minimap2` v2.17 [71]. Variants were called and a consensus sequence was reconstructed using `samtools/bcftools` v1.9 [72].

A phylogenetic tree was then reconstructed from the WGA with `RAxML` v8.2.12 [73] using the `GTRGAMMA` model and 1000 bootstrap replicates. The tree was rooted using midpoint.

Recombination analysis was performed from the same WGA using `Gubbins` v3.1.6 [74] with default settings. Figure 1 was generated using `Phandango` v1.3.0 [75]. The tree based on the 904 common genes of 61 *C. psittaci* strains (Figure S2) was constructed based on the core gene alignment produced by `RIBAP` and calculated using `RAxML` v8.2.12 [73].

A phylogenetic tree from SNP analysis was built using `RAxML` v8.2.12 with the `GTRGAMMA` model and 1000 bootstrap replicates based on the filtered SNP matrix (4011 SNPs) from `BioNumerics`.

### Multiple sequence alignments

Alignments of nucleotide and amino acid sequences were obtained using the `ClustalW` algorithm in `Geneious` 10.2.4 (Biomatters Ltd., Auckland, New Zealand). The `RAxML` tree based on OmpA sequences was constructed using protein model `GAMMA BLOSUM62` with the Rapid hillclimbing algorithm [73].

### Extraction of the PZ

The plasticity zone was identified by conducting a `BLAST` v2.10.0+ search against the gene *accB* as the 5’-terminus of the PZ and the gene *guaB* as the 3’-end. In case of a truncated PZ the MAC/perforin gene was taken as the 3’-end. Subsequently, genome fragments located within these boundaries were extracted using `bedtools v2.7.1 getfasta`.

### 3D structures of OmpA

OmpA protein structures were obtained using `ColabFold` [76], a free platform for prediction of 3D structures from amino acid sequences based on `AlphaFold` v2.1.0 [33, 77].

Graphics were generated using the molecular visualization program `ChimeraX` v1.4.dev202201150102, which was developed by the Resource for Biocomputing, Visualization, and Informatics at the University of California, San Francisco, with support from National Institutes of Health R01-GM129325 and the Office of Cyber Infrastructure and Computational Biology, National Institute of Allergy and Infectious Diseases [78].

### Multiple blast to identify homologs of Pmp and SinC proteins

All amino acid sequences of Pmp family members of *C. psittaci* strain 6BC were compiled in a multi-FASTA file and blasted against the proteome sequences of all 61 strains. The resulting hits were filtered to obtain those with the best bitscore for each individual Pmp in each strain. Likewise, the amino acid sequence of SinC of *C. psittaci* Cal10 (`EGF85279.1`) was blasted against all 61 proteomes.

### Statistical methods

To characterize possible relationships between isolation source/host and typing parameters of the *C. psittaci* strains, we assessed enrichment in a particular host using fisher exact test. For each combination of host and genotype, we assessed if a significant enrichment exists with respect to the proportions observed in the group of the remaining hosts.

## Data access

All used genomes including the reassembled ones, the `QUAST` and `IDEEL` quality results, as well as the complete core gene RIBAP output are deposited in an OSF repository: https://doi.org/10.17605/OSF.IO/RBCA9.

## Acknowledgements

We thank Hugues Richard, RKI MF1 Berlin, for his help with the statistical analysis.

## Footnotes

^1^ UniProt Feb 2022; available from: https://www.uniprot.org/uniprot/A0A656ISD9

^2^https://github.com/hoelzer/clean

^3^https://github.com/phiweger/ideel

^4^github.com/hoelzer-lab/ribap/tree/master/bin/helper/rearrange

^5^github.com/hoelzer-lab/ribap

^6^github.com/simonrharris/SKA

## Supplement

**Supplementary Figure S1:**
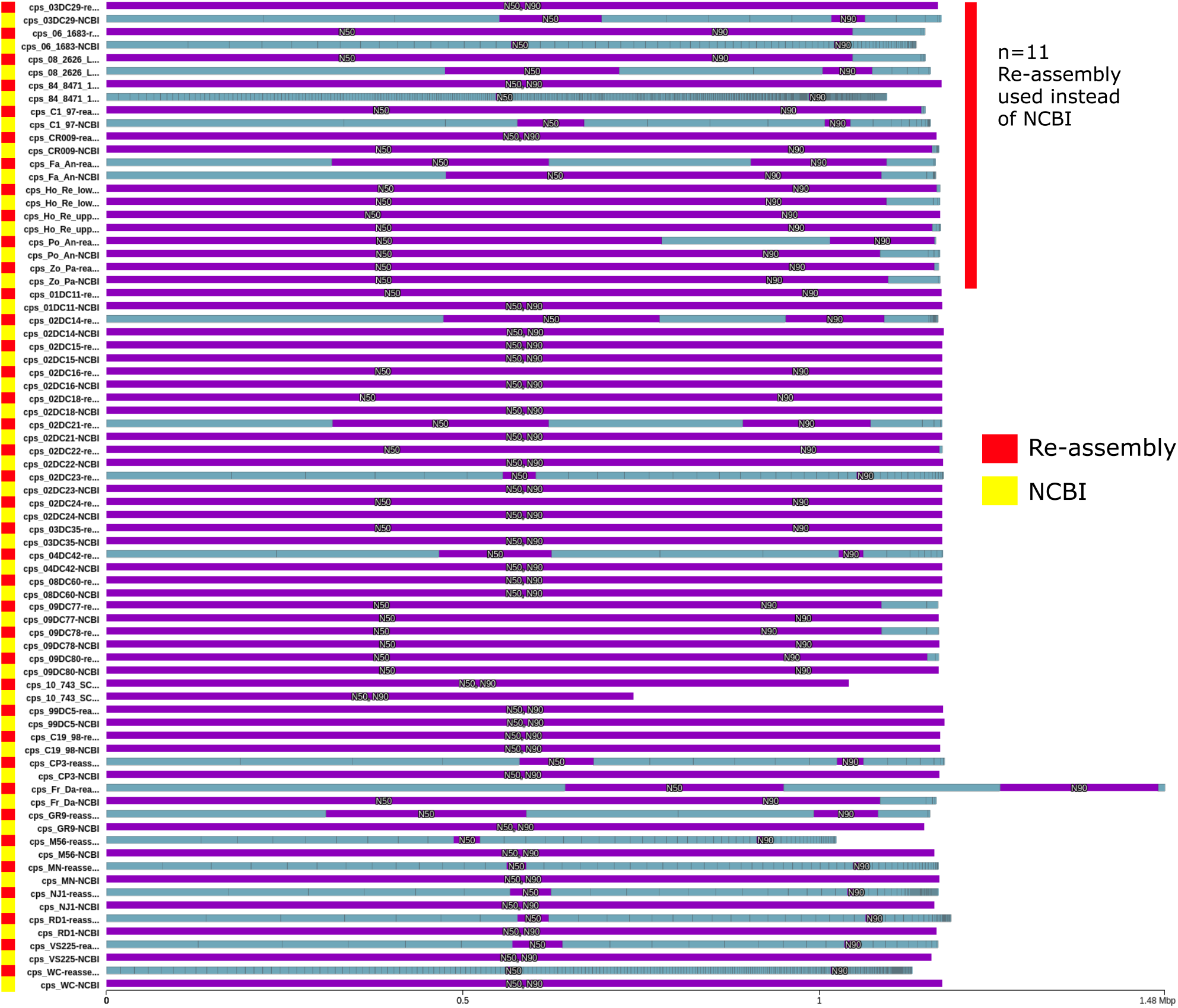
For *n* = 38 NCBI genomes we were able to find Illumina short-read sequencing data to reassemble these strains. The Icarus plot (output of `Quast v5.2.0` run with default parameters) shows the assembly contiguity and size for all 38 re-assemblies together with the corresponding genomes downloaded from NCBI. In addition, re-assemblies and NCBI genomes were decontaminated using CLEAN. The top eleven re-assemblies were finally selected to be integrated in our study and replaced the original NCBI genomes due to higher assembly contiguity and better N50 values. The purple bars mark contigs where a certain Nx (e. g. N50 or N90) is reached in an assembly.

**Supplementary Figure S2:**
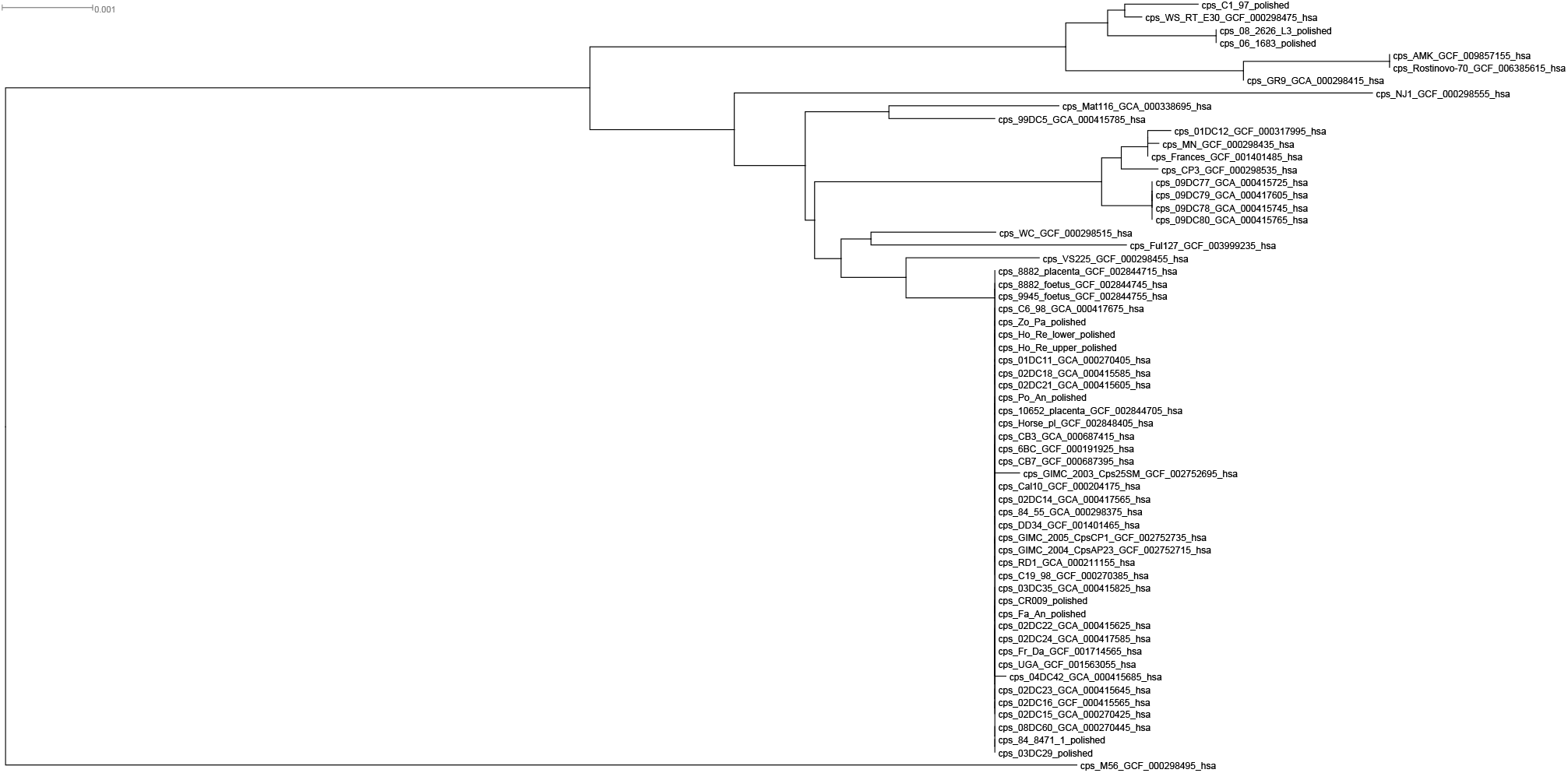
Phylogenetic tree based on the 904 common genes of 61 *C. psittaci* strains. The tree was reconstructed based on the concatenated core gene alignments at protein level produced by `RIBAP` and calculated using `RAxML` version 8.2.12 with the `GTRGAMMA` model and 1000 bootstrap replicates.

**Supplementary Figure S3:**
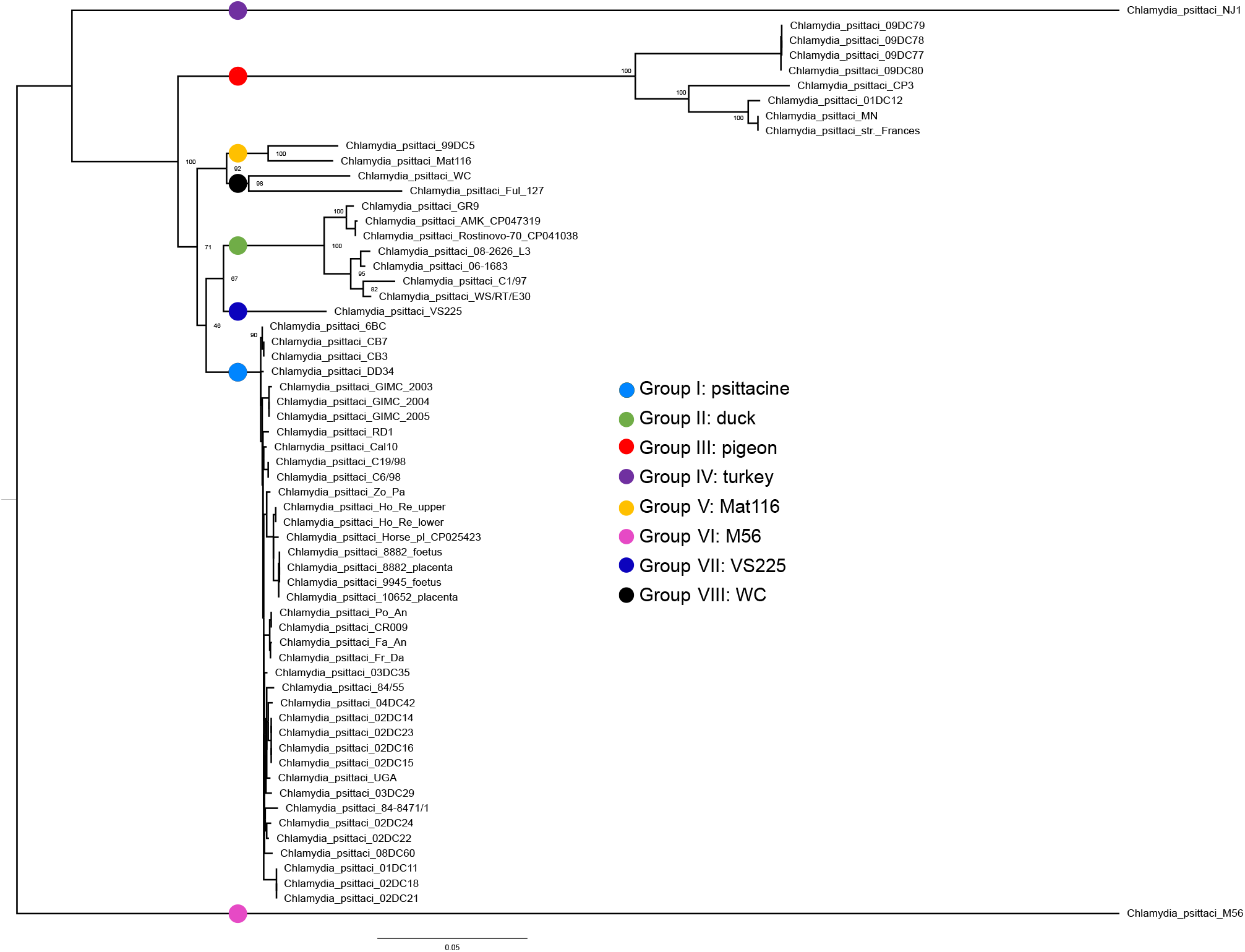
SNP-based tree determined from 61 *C. psittaci* genomes. The eight distinct lineages determined by Vorimore *et al.*, i.e. group I_psittacine, group II_duck, group III_pigeon, group IV_turkey, group V_Mat116, group VI_M56, group VII_VS225, and group VIII_WC are represented by colored circles. The tree was built using `RAxML` version 8.2.12 with the `GTRGAMMA` model and 1000 bootstrap replicates based on the filtered SNP matrix (4011 SNPs) from `BioNumerics`.

**Supplementary Figure S4:**
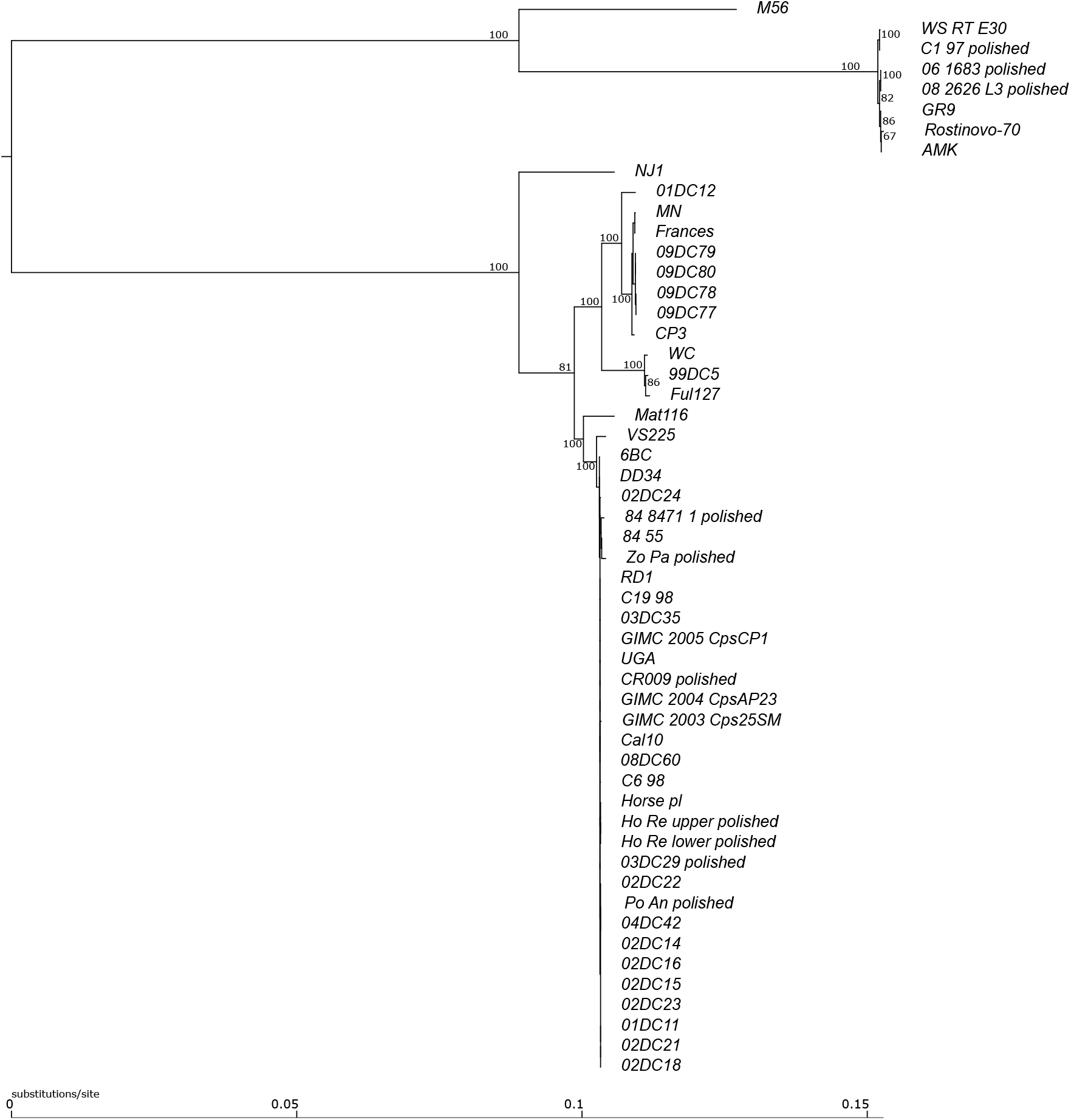
Phylogenetic tree based on nucleotide sequences of the extracted PZ of 53 *C. psittaci* strains used in this study. Sequences of 8 strains, where this region was located on several scaffolds, were not included here. The tree was constructed using `RAxML` v8.2.11 with `GTRGAMMA` nucleotide model and `Rapid hill-climbing` algorithm.

**Supplementary Figure S5:**
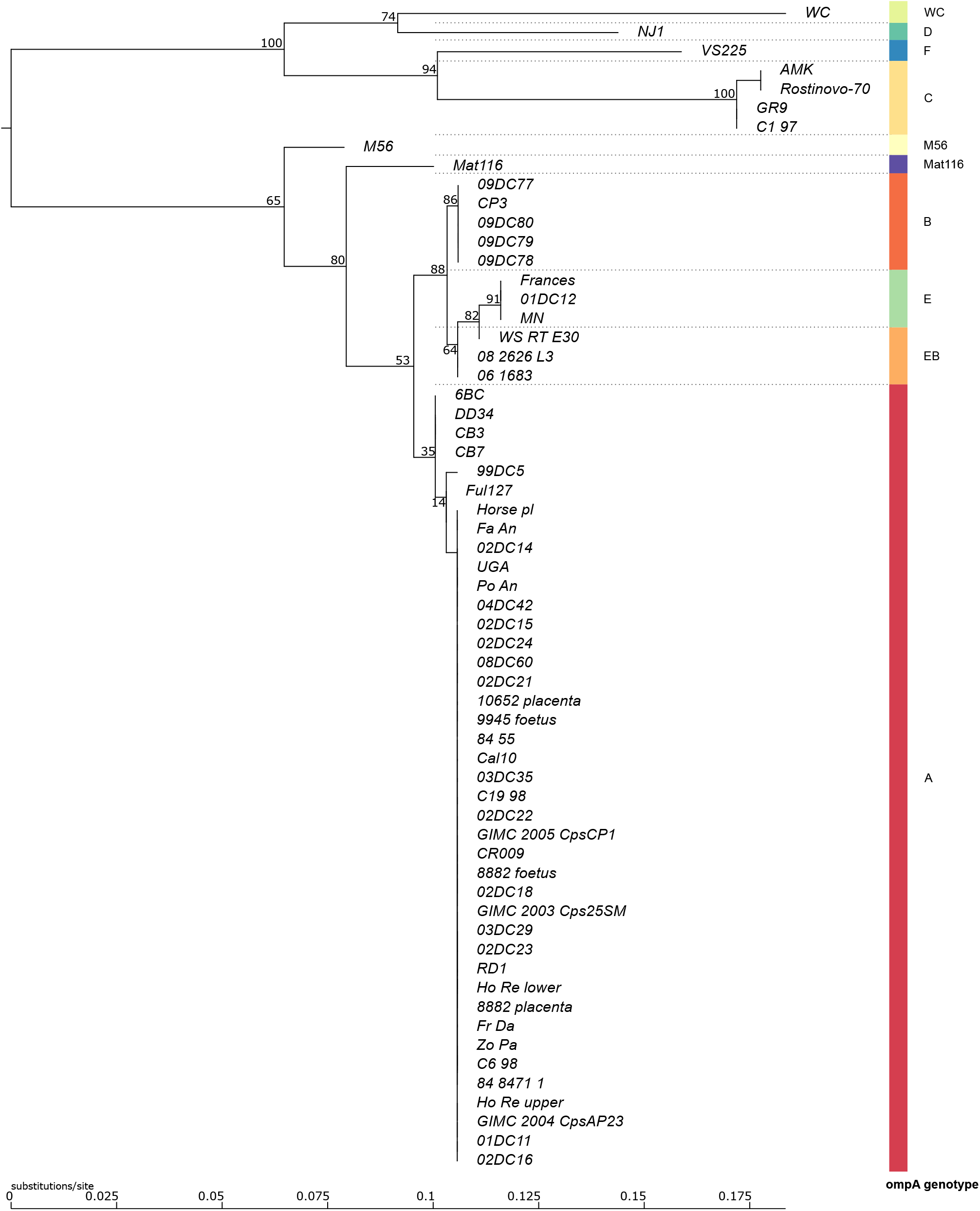
`RAxML` tree of the alignment of OmpA amino acid sequences from 61 *C. psittaci* strains as processed by `RIBAP` (Group 879). Bootstrap values are indicated at inner nodes. For identical taxa, bootstrap values were discarded, due to the interchangeability of corresponding gene sequences. The colored bar on the right indicates the respective *ompA* genotypes.

**Supplementary Figure S6:**
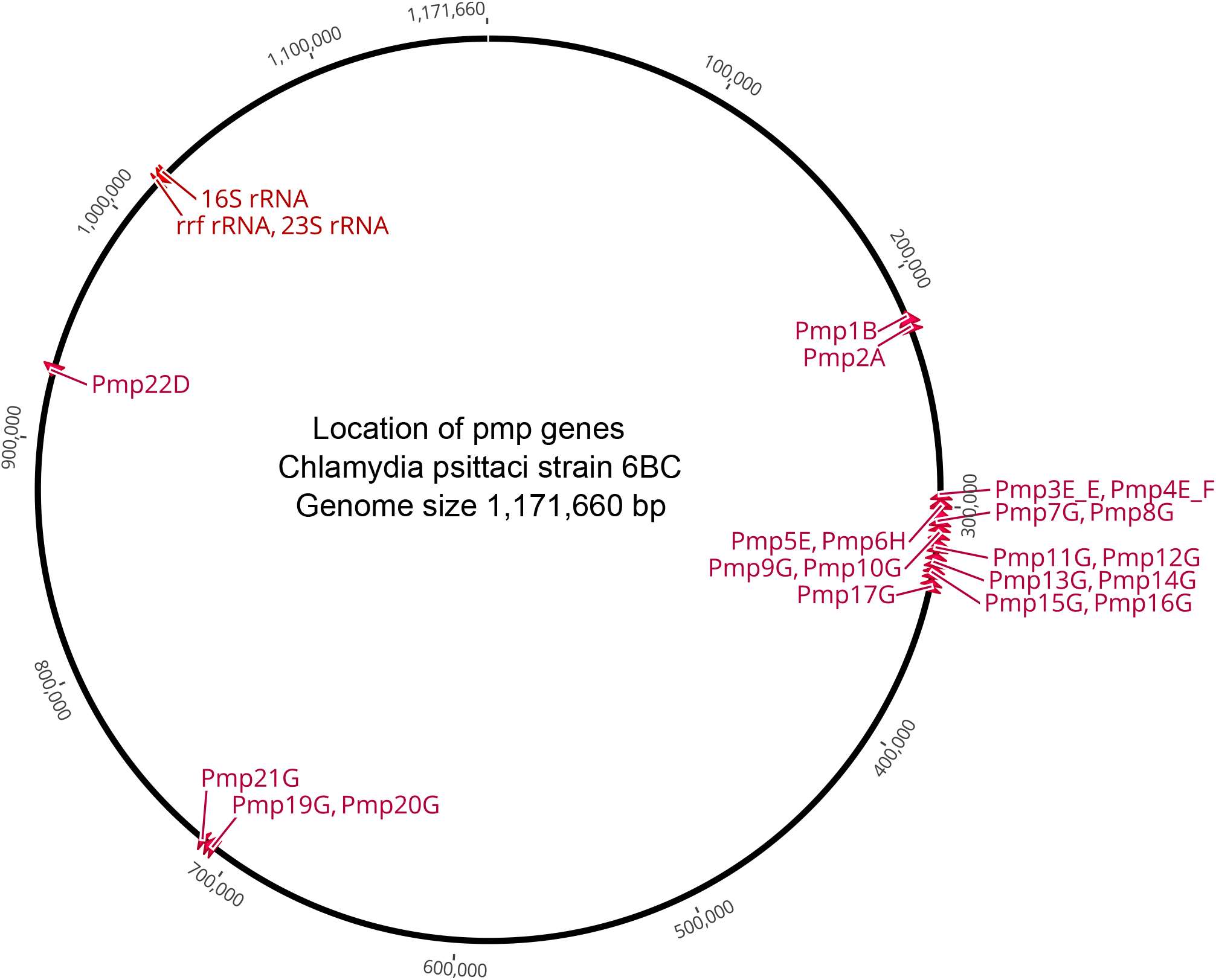
Location of genes encoding polymorphic membrane proteins in the genome of *C. psittaci* strain 6BC.

**Supplementary Table S1:** Basic characteristics, assembly state and typing data of all 61 strains included in this study. *incl. plasmids;**excl. plasmids;PZ 1* plasticity zone located on separate contigs.

**Supplementary Table S2:** Genetic distances (% identities) calculated from the nucleotide sequence alignment of complete plasticity zones of 53 *C. psittaci* strains.

**Supplementary Table S3:** Multiblast search for Pmp homologs to strain 6BC in 61 *C. psittaci* genomes (best hits).

**Supplementary Table S4:** Amino acid sequence variation among individual Pmps from 61 *C. psittaci* strains. *average aa identity of the respective Pmp to its 6BC homolog among all 61 strains

**Supplementary Table S5:** *C. psittaci* strains and host organisms used in this study.

